# Overlapping Pathogenic Signalling Pathways and Biomarkers in Preeclampsia and Cardiovascular Disease

**DOI:** 10.1101/2020.02.18.955260

**Authors:** Sonja Suvakov, Emma Bonner, Valentina Nikolic, Djurdja Jerotic, Tatjana P Simic, Vesna D Garovic, Guillermo Lopez-Campos, Lana McClements

## Abstract

**Objectives:** Preeclampsia is a cardiovascular pregnancy complication which occurs in 5-10% of pregnancies that can lead to a number of pregnancy complications including maternal and foetal death. Long-term, preeclampsia is associated with up to 8-fold increased risk of cardiovascular disease (CVD) for both mothers and their offspring. The lack of mechanistic data in relation to the causes or consequences of preeclampsia has prevented the development of effective therapeutic or monitoring strategies.

**Study design:** This study investigates common underlying mechanisms of preeclampsia and CVD, specifically hypertension and heart failure with preserved ejection fraction (HFpEF) using “*in silico”* approach of publicly available datasets. Integrated techniques were designed to mine data repositories and identify relevant biomarkers associated with these three conditions.

**Main outcomes measures:** The knowledge base tools were employed that enabled the analysis of these biomarkers to discover potential molecular and biological links between these three conditions.

**Results:** Our bioinformatics “*in silico*” analyses of the publically available datasets identified 76 common biomarkers between preeclampsia, hypertension and HFpEF. These biomarkers were representative of 29 pathways commonly enriched across the three conditions which were largely related to inflammation, metabolism, angiogenesis, remodelling, haemostasis, apoptosis, endoplasmic reticulum (ER) stress signalling and the renin-angiotensin-aldosterone (RAAS) system.

**Conclusions:** This bioinformatics approach which uses the wealth of scientific data available in public repositories can be helpful to gain a deeper understanding of the overlapping pathogenic mechanisms of associated diseases, which could be explored as biomarkers or targets to prevent long-term cardiovascular complications such as hypertension and HFpEF following preeclampsia.

**Highlights:**

- Women with preeclampsia have increased risk of cardiovascular disease later in life but the mechanism is poorly understood.
- “*In silico*” analyses of publically available datasets provided overlapping biomarkers and pathogenic pathways between preeclampsia, hypertension and heart failure with preserved ejection fraction (HFpEF).
- These data could be utilised in the future studies that may lead to the development of better risk stratification strategies or preventative treatments for women post preeclampsia.

## Introduction

Preeclampsia is a cardiovascular complication of pregnancy affecting 5-10% of all pregnancies annually. It is clinically characterised by the new onset hypertension (≥140/90mmHg) and the presence of proteinuria (≥300mg/day) or other organ damage that develops after 20 weeks of gestation with limited treatment options (1). Preeclampsia is a multifactorial condition with molecular regulation of its pathogenesis remaining poorly understood. However, the development of preeclampsia has been closely associated with abnormal placentation occurring in the early stages of pregnancy where impaired spiral uterine artery (SUA) remodelling plays a significant role(2). The lack of the SUA remodelling represents an initial step leading to vascular changes and hypertension(3).

Longitudinal studies reported that endothelial dysfunction in women whose pregnancy was complicated by preeclampsia persists for 10-20 years post-partum, particularly following severe preeclampsia cases (4). Preeclampsia is associated with up to an 8-fold increased risk of CVD in later life for both mothers and their offspring (5). Although it is unknown whether pre-existing cardiovascular risks (CVRs) become uncovered due to cardiovascular stresses of pregnancy or if the endothelial damage caused by preeclampsia increases the CVR, there remains a strong link between preeclampsia and the development of CVD in later life. CVD in women is the global leading cause of female death (6). Preeclampsia and hypertension are well-known risk factors for the development of heart failure, more commonly associated with heart failure with preserved ejection fraction (HFpEF)(7,8). HFpEF is an asymptomatic condition clinically characterised by diastolic dysfunction that results in reduced ventricular relaxation with preserved contractile function, maintaining an ejection fraction of ≥50% (9), and it is more common in women than men (10).

The wealth of knowledge of human disease is continuously growing due to daily publications on data repositories such as the National Centre for Biotechnology Information (NCBI). Various tools have been developed to mine and retrieve data from biomedical literature in relation to the expression levels of certain biomarkers within specific diseases and how they interact to modify the function of various pathways(11). Due to the immense volume of biomedical literature available on NCBI, MeSH terms were introduced to improve the literature search and retrieval process (12). These allow for the retrieval of documents that possess similar content but use various synonyms, abbreviations or wording regarding the same biological term e.g. ‘preeclampsia’ or ‘pre-eclampsia,’ allowing for the construction of a refined and specific literature search (12). Reactome, a data analysis tool, which contains 10719 human genes interacting into 2102 pathways, was designed to aid in the discovery of biological interactions and reactions occurring under normal physiological and disease conditions (13).

We hypothesise that there are common underlying pathways involved in the pathogenesis of both preeclampsia and CVD, specifically hypertension and HFpEF. To test this hypothesis, an “*in silico”* approach was applied which investigated biomedical literature provided by public repositories to analyse the biomarkers differentially expressed in preeclampsia, hypertension and HFpEF and to identify overlapping biological pathways consistent across all three conditions.

## Materials and Methods

All the analyses were implemented as an R script using 3.6.0. This script contained the following sections:

### Literature search and retrieval

The literature was searched and retrieved using NCBI relevant to preeclampsia, hypertension and HFpEF using various PubMed queries:

- The preeclampsia query was structured as ‘((“pre-eclampsia”[MeSH Terms] OR “pre-eclampsia”[TIAB] OR “preeclampsia”[TIAB]) AND (biomarkers”[MeSH Terms] OR “biomarkers”[TIAB] OR “biomarker”[TIAB]) AND (“1900/01/01”[Date - Entrez]: “2019/06/01”[Date - Entrez]))’ - the query aimed to retrieve documents containing specific biomarkers of preeclampsia.
- The hypertension query was structured as ‘((Hypertension [MeSH] NOT preeclampsia) AND (“biomarkers”[MeSH Terms] OR “biomarkers”[TIAB] OR “biomarker”[TIAB]) AND (“1900/01/01”[Date - Entrez]: “2019/06/01”[Date - Entrez]))’ - this query aimed to retrieve documents annotated for hypertension while excluding preeclampsia as a hypertensive state.
- The HFpEF query was structured as ‘((HFpEF [TIAB] OR “heart failure with preserved ejection fraction “[TIAB]) AND (“biomarkers”[MeSH Terms] OR “biomarkers”[TIAB] OR “biomarker”[TIAB]) NOT preeclampsia AND “1900/01/01”[Date - Entrez]: “2019/06/01”[Date - Entrez]))’ - this query aimed to retrieve documents annotated with biomarkers for HFpEF.

The R package RISMed (2.1.7) package was used to download the contents from the retrieved literature into R. To identify the relevant documents in PubMed, the queries were passed as arguments to the “EUtilsSummary” function. These documents were downloaded into R by the “EUtilsGet” function constructing three data frames containing the PubMed ID, NCBI gene ID and gene name for each biomarker retrieved from the queries.

### Biomarker Annotation and Extraction

Pubtator, a tool designed to annotate biological terms cited in PubMed documents, was used to annotate the biomarkers cited within the literature retrieved from the three queries (14). The PubMed IDs obtained from the literature search and retrieval were used as input into Pubtator as previously described (11).

This single gene ID file was read into R using the “read.csv” function. The Jaccard index was calculated using the “Jaccard” function to indicate the similarity between the gene sets. This function compared the similarity between preeclampsia and hypertension, preeclampsia and HFpEF and hypertension and HFpEF. From this, the Jaccard distances were calculated, illustrating the level of dissimilarity between the gene sets. The Jaccard distance is defined as 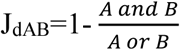, where A and B are distinct sets (15).

### Gene Set Analyses

Three gene set analyses were conducted to identify which pathways the biomarkers identified in preeclampsia, hypertension and HFpEF, are a part of, and determine if any pathways are significantly enriched across all three conditions using Reactome knowledgebase. The enrichment analyses applied a hypergeometric test to assess whether the number of genes associated with a reactome pathway was larger than would be expected at random. These tests were calculated using the NCBI Gene Identifiers obtained from the literature analyses as input for the the R package ReactomePA, for the analyses using reactome database, and were corrected for multiple comparisons using a “Benjamini-Hochberg” correction. Statistical significance threshold was set to adjusted p-value less than 0.05 (adj.p-value <0.05).

## Results and Discussion

### Retrieval and Annotation of Biomedical Literature with Biomarkers

A total of 3,007 biomarkers were annotated across the 5,738 documents, with 76 biomarkers being present in all three gene sets, indicating only a 2.5% overlap between preeclampsia, hypertension and HFpEF (Fig. 1). Within the preeclampsia datasets, 779 of the biomarkers annotated were unique to preeclampsia, whereas 1,188 were unique to hypertension and 30 were unique to HFpEF. A total of 462 biomarkers overlapped between preeclampsia and hypertension (25 % overlap), 84 overlapped between preeclampsia and HFpEF (6.7% overlap) and 110 overlapped between hypertension and HFpEF (6.5% overlap). A Jaccard index of approximately 0.016 was found between hypertension and HFpEF, generating a Jaccard distance of 0.984. Similar findings were discovered for the other gene sets with a Jaccard index of 0.016 and distance of 0.984 reported between preeclampsia and HFpEF and an index of 0.04 and distance of 0.96 reported between preeclampsia and hypertension. These figures illustrate a high level of dissimilarity between the preeclampsia, hypertension and HFpEF gene sets.

**Figure 1.**
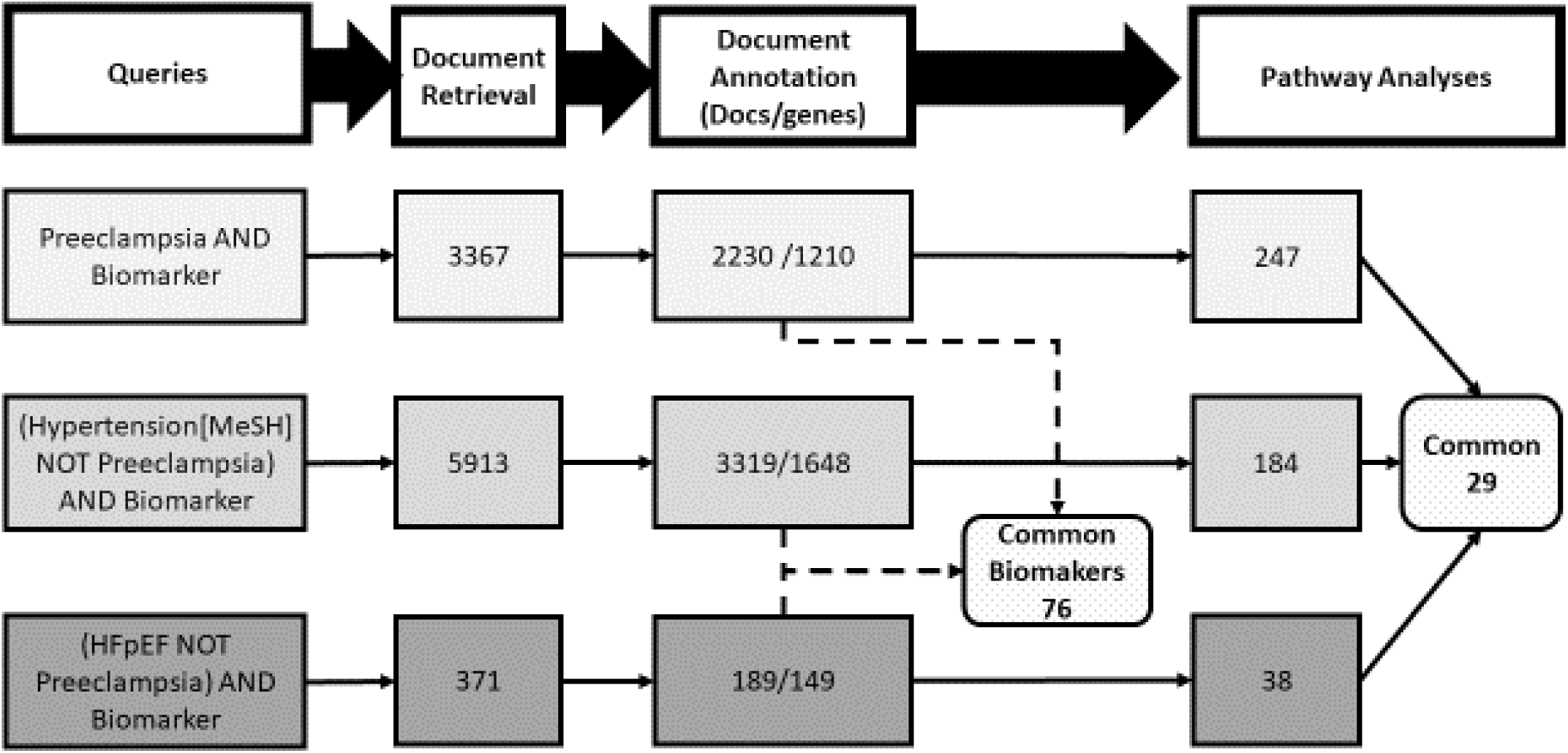
A schematic of data mining and analysis bioinformatics strategies used and the results obtained. This figure illustrates a summary of the methods used and the results obtained based on the retrieval of the documents annotated with biomarkers and the number of pathways found to be significantly enriched across all three disease states by a Reactome analysis of the gene sets.

### Identifying overlapping pathways between preeclampsia, hypertension and HFpEF

Once the biomarkers identified in these three conditions were determined, Reactome was used to extrapolate these findings into enriched pathways potentially involved in the pathogenesis of preeclampsia, hypertension and HFpEF. The preeclampsia gene set was most significantly enriched with 247 pathways, hypertension with 184 and HFpEF with 38; 29 pathways were enriched in all three gene sets (Figure 2A). A fourth gene set analysis of the 76 common biomarkers discovered 31 significantly enriched pathways in this gene set. Out of the 295 pathways significantly enriched across the four gene sets, 25 were commonly enriched in all four gene sets (Figure 2B). The pathways enriched between the three conditions are presented in Figure 3. Specific pathogenic pathways identified for each condition are presented in Supplementary Fig. 1-3).

**Figure 2.**
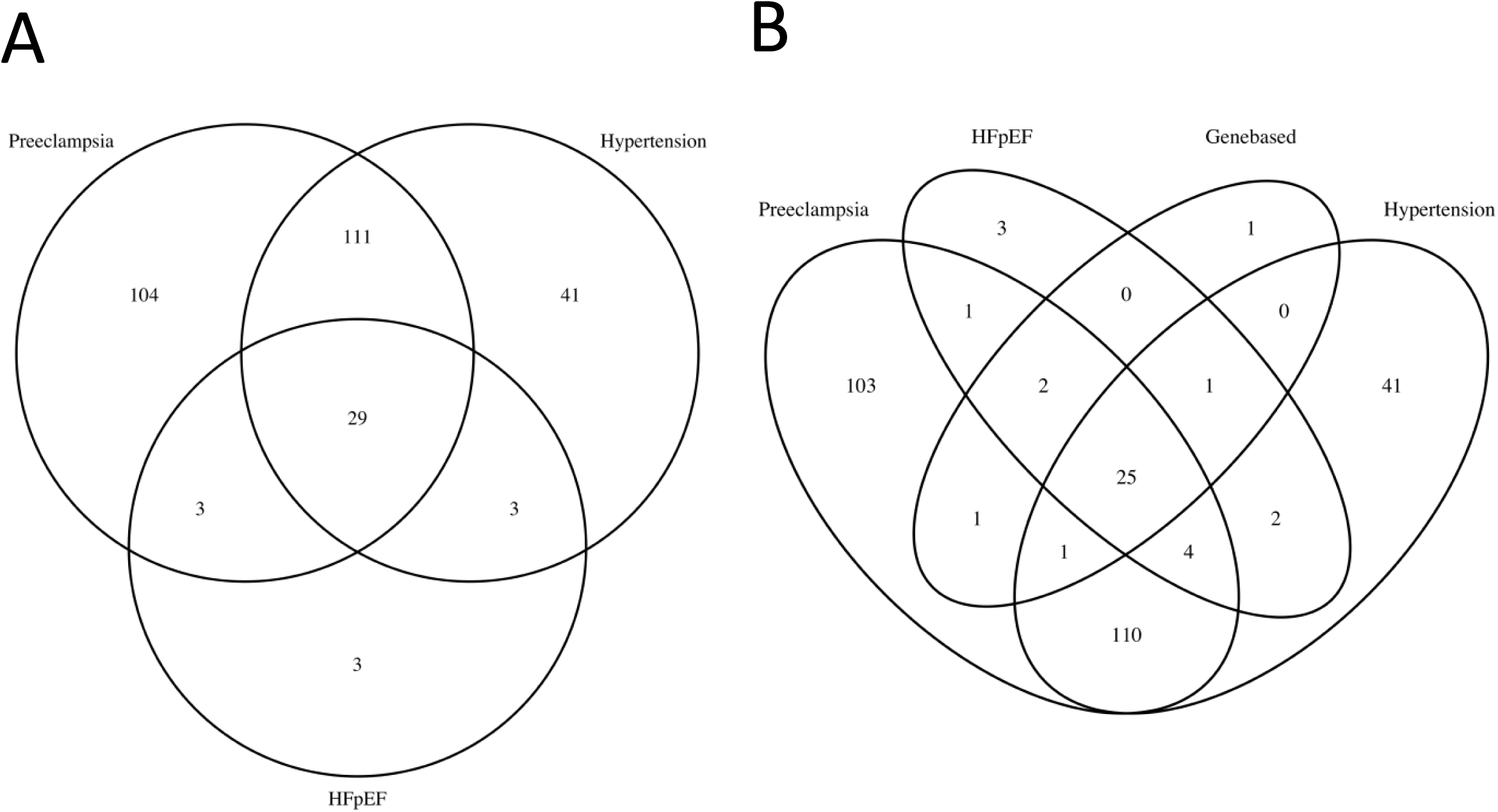
Gene set analysis showing the significantly enriched pathways in preeclampsia, hypertension and HFpEF. An analysis of the annotated biomarkers on Reactome found 247 pathways were significantly enriched in preeclampsia, 104 of which were unique to preeclampsia, 183 in hypertension with 41 pathways being unique to this condition and 38 in HFpEF, 3 of which were unique to HFpEF. Of the 291 statistically significant pathways that were enriched across all three conditions, 29 of these were found to be common in preeclampsia, hypertension and HFpEF **(Figure 2A)**. A fourth gene set analysis including the 75 common biomarkers in preeclampsia, hypertension and HFpEF found that 31 pathways were significantly enriched in this gene set. An overlap analysis of the 295 significantly enriched pathways across the four gene sets found 29 pathways were commonly enriched in each gene set **(Figure 2B).** Significance was determined by a threshold of an adjusted p-value<0.05. Abbreviations: HFpEF=heart failure with preserved ejection fraction.

**Figure 3.**
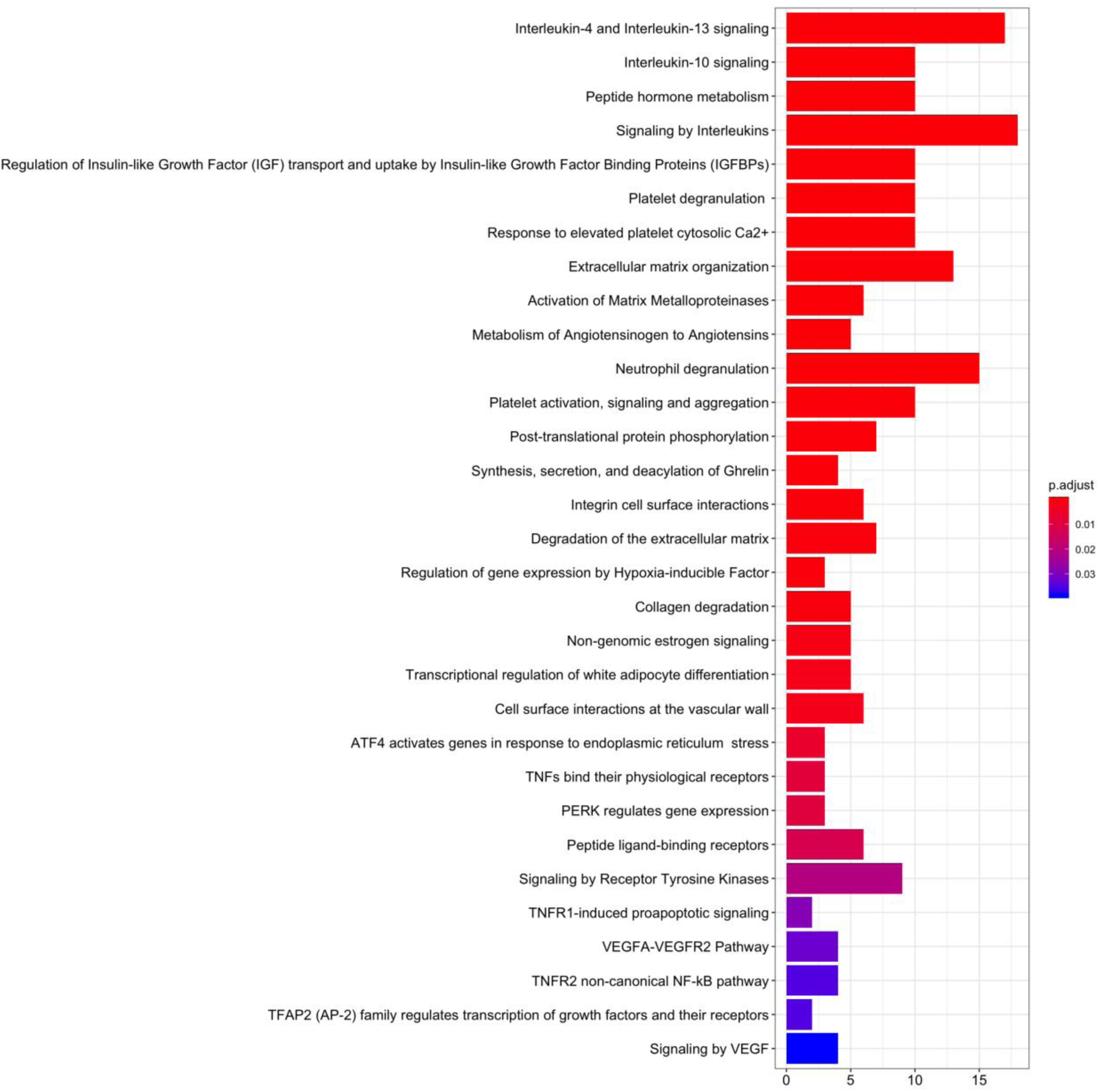
Significantly overlapping signalling pathways between preeclampsia, hypertension and heart failure with preserved ejection fraction. Reactome knowledge base was used to identify overlapping pathways between preeclampsia, hypertension and heart failure with preserved ejection fraction (HFpEF) based on the common genes/biomarkers identified in the initial analysis.

### Overlapping pathophysiological mechanisms between preeclampsia, hypertension and HFpEF

The bioinformatics analyses of publically available datasets identified 76 commonly expressed biomarkers in preeclampsia, hypertension and HFpEF. Preeclampsia datasets showed the highest pathway enrichment which reflect the complex pathogenesis of preeclampsia at the molecular level.

Considering that women who experience preeclampsia during pregnancy have higher burden of cardiovascular complications later in life (16), in this study we identified a number of overlapping molecular pathways between preeclampsia and CVD i.e. hypertension and HFpEF. These include pathways associated with metabolism, angiogenesis, vascular remodelling, haemostasis, apoptosis and ER signalling (Figure 3). The increasing body of evidence supports the immune system activation and inflammation as crucial steps involved in pathogenesis of preeclampsia(17) hypertension (18), and HFpEF (19). Existing data show pro-inflammatory cytokines and mediators such as interleukin-6 (IL-6), IL-1, IL-8, C-reactive protein (CRP), monocyte chemoattractant protein-1 (MCP-1) and tumour necrosis factor-alpha (TNF-α) to be overexpressed in preeclampsia, hypertension and HFpEF, suggesting inflammation as an important underlying mechanism(20,21). A study showed that within reported CRP concentrations in patients with coronary artery disease, this well-known marker of inflammation is able to induce endothelial dysfunction by decreasing endothelial nitric oxide synthase activity (22). Interestingly, markers of immune response, such as CD4+ and CD8+ T-cells were also annotated in the preeclampsia and hypertension gene sets, whereas not in HFpEF gene set (21)-(23). Both CD4+ and CD8+ T-cells are dysregulated in these diseases and produce pro-inflammatory cytokines which can provoke or maintain high blood pressure (24,25). In line with other pro-inflammatory molecules, IL6 and TNF-alpha have been associated with structural and functional changes in endothelial cells (26). On the other hand, anti-inflammatory cytokine such as IL-10 was reported to be downregulated in all three diseases (21). Decreased production of IL-10 leads to vascular dysfunction in association with increased endothelin-1 and further causes an increase in pro-inflammatory cytokines such as IL6 and TNF-α possibly through the ERK1/2 pathway(27). In our analysis, endothelin-1, IL-10 and TNF-α as well as other inflammatory cytokines and chemokines (IL1A, IL6, CXCL8) were implicated in all three conditions.

ER stress and apoptosis signalling pathways including PERK/AFT4, TNFR1/TNFR2 and tyrosine kinases were also identified in our study as overlapping mechanisms between these three conditions, suggesting that these pathways could represent potential therapeutic targets for pre-eclampsia, hypertension and HFpEF. There are already available therapeutics used for the treatment of different inflammatory conditions, which target some of the substrates in these pathways(28).

Although endothelin-1 and cyclooxygenase-dependent vasoconstrictors may have the predominant role in the development of hypertension, many studies have demonstrated an important role of oxidative stress in hypertension(29,30) which aligns to our findings. Notably, NADPH oxidases (NOX4 and NOX5) was found to be annotated in this study in both hypertension and preeclampsia but not in HFpEF. This could have an impact on the production of superoxide anion which determines biological availability of NO, since these two molecules bind and form peroxynitrites, ONOO- (31). Another link of reactive oxygen species production and NO availability is found in the processes of eNOS uncoupling. Namely, when tetrahydrobiopterin (BH4) levels are diminished, due to decreased activity of a rate limiting enzyme for BH4 production - GTP cyclohydrolase, it results in lower NO synthesis and enhanced superoxide anion production (32). However, depletion of BH4 could also be a result of its catabolism as a result of peroxynitrite attack indicating that oxidative stress is one of the underlining mechanisms of endothelial dysfunction (33). In our study, the enrichment of biomarkers involved in angiogenesis and vascular remodelling included TGF-β, Gal-3, VEGF, endoglin, collagen group proteins and MMPs were also implicated in all three conditions. A well-known compensatory pro-angiogenic factor activated by hypoxia(34), hypoxia-inducible factor 1α (HIF-1α) was also identified in all three conditions. These findings align with well-established knowledge that one of the root causes of preeclampsia is inappropriate implantation and development of the placenta due to poor SUA remodelling (2,35). Hypertension and HFpEF are also characterised by aberrant angiogenesis, vascular remodelling and fibrosis(36,37). Furthermore, von Willebrand Factor (vWF) and fibrinogen, biomarkers annotated in all three gene sets, are essential for the formation of stable blood clots, therefore playing an important role in haemostasis in preeclampsia, hypertension and HFpEF (38). In addition to their important role in haemostasis, vWF represents a plasma marker of endothelial damage(38,39). Inverse association between vWF and flow-mediated vasodilatation (40) suggests that endothelial NO levels may have a negative feedback on vWF secretion. It was also demonstrated that endothelial secretion of vWF is mediated by several inflammatory cytokines(41). In our study we also identified selectin-E and endothelin-1 as overlapping biomarkers between the three conditions; all of which are correlated with endothelial dysfunction(42,43). There is conflicting evidence surrounding the overlap of biomarkers expressed in preeclampsia and HFpEF. Alma *et al* found CRP, high density lipoprotein (HDL), insulin, fatty acid binding protein-4 (FABP4), brain natriuretic peptide (BNP), N-Terminal pro-BNP (NT-proBNP), adrenomedullin (ADM), mid-regional pro-ADM, cardiac Troponin-1 (c-Tn1) and cancer antigen-125 (CA-125) as commonly expressed in preeclampsia and HFpEF (44). Some of these biomarkers such as BNP/NT-proBNP, CRP, CA-125, insulin and c-Tn1 were also identified in our analyses as overlapping between the three conditions investigated. Biomarkers such as ADM/MR-proADM, HDL and FABP4 were not annotated for HFpEF.

A number of overlapping metabolic pathways were identified including peptide hormone metabolism, insulin-like growth factor (IGF) transport and IGF binding proteins (IGFBP), and transcriptional regulation of white adipocyte differentiation. These pathways were determined in association with the following biomarkers: dipeptidyl peptidase 4, insulin like growth factor 1, IGFBP-1, insulin, lipocalin 2, leptin, Gal-3. It is well-established that obesity and diabetes are associated with increased risk of pre-eclampsia in pregnancy(45), hypertension(46) as well as HFpEF where diabetes and obesity are linked to poorer outcomes for the patients(47). Metabolism by renin of angiotensinogen to angiotensin was also identified as one of the significant pathways in all three conditions which could potentially represent an effective preventative or treatment strategy during and post pre-eclampsia. Nevertheless, the safety of the therapeutics inhibiting the RAAS system in pregnancy is either lacking or found unsafe. Previously published literature has reported a positive association between blood glucose and the aldosterone levels as well as a negative correlation between the plasma renin and blood glucose levels suggesting that the RAAS system might be implicated in insulin resistance or new onset diabetes in pregnancy(48).

Although preeclampsia is a distinct cardiovascular pregnancy complication associated with increased risk of cardiovascular disease in the future, there are substantial differences in the molecular regulation of the pathogenesis of preeclampsia compared to hypertension or HfpEF with some shared common pathways, including inadequate immune response, inflammation, vascular homeostasis and ER stress.

### Limitations and Future Work

The main limitations of this research surround the methodologies used to obtain the literature and datasets used for the analysis of the biomarkers expressed in preeclampsia, hypertension and HFpEF. The annotation process which used Pubtator to mine the biomedical literature assumed no errors were made in the annotation of the biomarkers included in the gene sets and did not acknowledge a potential for the inclusion of biomarkers that were falsely identified (false positives), or biomarkers that were missed and not included in the gene sets (false negatives). Also our analysis assumed that all biomarkers cited within the literature were associated with the disease.

## Conclusion

There are limited existing studies thus far that have researched the possibility of an underlying common pathogenesis between preeclampsia, hypertension and HFpEF. Our “*in silico”* approach enabled the identification of 76 common biomarkers between the three conditions that were predominantly involved in pathways related to metabolism, inflammation, vascular remodelling, angiogenesis, haemostasis, ER stress signalling and the RAAS system. Therefore, this could be basis for future research to further investigate viable biomarkers or targets in order to identify a potential treatment or monitoring strategy to reduce the global morbidity and mortality rates associated with preeclampsia and subsequent CVD.

## Funding

This work was supported by the Faculty of Science Research and Development Fund at University of Technology Sydney.

## Detail of ethics approval

No ethical approval was required.

## Declaration of interests

No conflicts of interest are declared.

---

**Supplementary Figure 1**. Pathogenic pathways identified by Reactome in heart failure with preserved ejection fraction (HFpEF).

**Supplementary Figure 2**. Pathogenic pathways identified by Reactome in hypertension (HT).

**Supplementary Figure 3**. Pathogenic pathways identified by Reactome in preeclampsia (PE).

